# Efficient minimizer orders for large values of *k* using minimum decycling sets

**DOI:** 10.1101/2022.10.18.512682

**Authors:** David Pellow, Lianrong Pu, Baris Ekim, Lior Kotlar, Bonnie Berger, Ron Shamir, Yaron Orenstein

## Abstract

Minimizers are ubiquitously used in data structures and algorithms for efficient searching, mapping, and indexing of high-throughput DNA sequencing data. Minimizer schemes select a minimum *k*-mer in every *L*-long sub-sequence of the target sequence, where minimality is with respect to a predefined *k*-mer order. Commonly used minimizer orders select more *k*-mers than necessary and therefore provide limited improvement in runtime and memory usage of downstream analysis tasks. The recently introduced universal *k*-mer hitting sets produce minimizer orders with fewer selected *k*-mers. Unfortunately, generating compact universal *k*-mer hitting sets is currently infeasible for *k* > 13, and thus cannot help in the many applications that require minimizer orders for larger *k*.

Here, we close the gap of efficient minimizer orders for large values of *k* by introducing *decycling-set-based minimizer orders*, new orders based on minimum decycling sets. We show that in practice these new minimizer orders select a number of *k*-mers comparable to that of minimizer orders based on universal *k*-mer hitting sets, and can also scale up to larger *k*. Furthermore, we developed a method that computes the minimizers in a sequence on the fly without keeping the *k*-mers of a decycling set in memory. This enables the use of these minimizer orders for any value of *k*. We expect the new orders to improve the runtime and memory usage of algorithms and data structures in high-throughput DNA sequencing analysis.

## 1 Introduction

As the number and depth of high-throughput sequencing experiments grows, efficient methods to map, store, and search DNA sequences have become critical in their analysis. Sequence sketching is a fundamental building block of many of the basic sequence analysis tasks, such as assembly (Rautiainen and Marschall, 2021; Ekim et al., 2021), alignment (Sahlin, 2022; Dutta et al., 2022; Li, 2018), binning (Deorowicz et al., 2015; Chikhi et al., 2016; Flomin et al., 2022), and indexing (Marchet et al., 2021; Holley and Melsted, 2020). The common principle in all sketching techniques is the consistent selection of representative *k*-mers from longer DNA sequences for indexing these sequences in data structures or algorithms. A key parameter for evaluating and comparing sketching schemes is density (Marçais et al., 2017), which is defined as the fraction of *k*-mers selected from a sequence by the scheme.

One of the most common sequence sketching techniques is minimizers (Schleimer et al., 2003). The minimizer of an *L*-long sequence is the minimum among all the *k*-mers that the sequence contains, according to some order o over the *k*-mers. Selecting the minimizers from all *L*-long windows of a sequence provides a sketch of that sequence. Minimizer schemes provide a window guarantee, i.e., one representative *k*-mer is selected from every *L*-long window for any desired value of L. Minimizers have low density as the minimum *k*-mer is likely to persist across multiple overlapping windows. However, the commonly used lexicographic and random *k*-mer orders have been shown to have far from optimal density (Marçais et al., 2017).

A recent breakthrough in developing minimizer orders with low density has been achieved by compact universal *k*-mer hitting sets (UHSs) (Orenstein et al., 2017). A UHS is a set of *k*-mers guaranteed to hit (i.e., have a member included in) any *L*-long sequence. In terms of a complete de Bruijn graph (dBG) of order *k*, a minimum UHS is a minimum set of nodes whose removal leaves no path of length *L* – *k* +1 in the graph. Heuristic algorithms for finding a compact UHS include DOCKS (Orenstein et al., 2017) and PASHA (Ekim et al., 2020), both of which approach UHS construction as a path covering problem in a complete dBG. Both algorithms first identify a minimum decycling set (MDS), which is a set of *k*-mers guaranteed to hit any infinitely long sequence, and then extend this set into a UHS. An MDS can be generated in time linear in the dBG size (Mykkeltveit, 1972).

UHS-based minimizer orders were shown to achieve lower density than common orders (Marçais et al., 2017; Ekim et al., 2020). However, constructing and storing UHSs is inefficient due to the exponential dependence of the heuristic algorithms on *k*, and currently compact UHSs are available only for *k* ≤ 13. As a result, UHS-based minimizer orders could not be utilized in many applications that require larger values of *k*, such as long-read mapping (Li, 2018), assembly (Rautiainen and Marschall, 2021), and indexing (Holley and Melsted, 2020; Pibiri, 2022). Moreover, another bottleneck arising in these applications is storing and querying UHS *k*-mers, as the UHS size grows exponentially with *k*.

Partly due to the challenges in constructing UHSs, other recent works focused on developing sequence-specific minimizer orders. For example, sequence-specific minimizer orders were used in binning applications to achieve lower maximum bin size or more balanced bin sizes than general minimizers (Chikhi et al., 2016; Flomin et al., 2022). Hoang et al. (2022) used deep learning to achieve sequence-specific low-density minimizers for much larger *k* (up to 320). Polar-set-based sequence-specific minimizers achieved very low density for *k* ≤ 25 (Zheng et al., 2021). These solutions are tailored to a specific sequence set, and thus different orders must be generated for every sequence.

In this work, we developed new methods to construct universal, low-density minimizer orders that scale to larger *k*. Motivated by the fact that current algorithms, DOCKS and PASHA, generate a UHS on top of an MDS, we defined minimizer orders based only on an MDS. For most combinations of *k* and *L*, most of the *k*-mers in the UHSs generated by these algorithms are in the MDS. In addition, generating an MDS is the fastest part of UHS construction in these algorithms, taking only *Θ*(|*Σ*|*^k^*) for alphabet *Σ*. Moreover, the same order is used for all *L* and a given *k*, unlike DOCKS and PASHA, which need to compute a UHS for every combination of *k* and L. Furthermore, the longest remaining path in a complete dBG after removal of an MDS was recently shown to be bounded by *O*(*k*^3^) (Zheng et al., 2020a), so for large values of *L* it is likely that many of the windows contain a *k*-mer from an MDS.

As the MDS size grows exponentially in *k*, pre-computing, storing, and querying it becomes infeasible for large *k*. We overcome this limitation by presenting an efficient method to test in *O*(*k*) time if a *k*-mer belongs to the MDS, which enables computing the minimizers in a sequence according to our MDS-based order on the fly. We demonstrate that our new MDS-based minimizer orders achieve density that is comparable to or better than UHS-based orders. The minimizer orders we defined thus provide, for the first time, universal orders with low density that easily scale to any value of *k*, and any desired window length. All code developed under this project is publicly available via github.com/OrensteinLab/DecyclingSetBasedMinimizerOrder.

## 2 Preliminaries

We begin by providing definitions and theoretical background and describing relevant related work. See (Orenstein et al., 2017; Edgar, 2021) for further background.

### 2.1 Basic definitions

For a string *S* over an alphabet *Σ*, a *k-mer* is a contiguous substring of length *k*. We denote the *k*-mer starting at position *i* as *S*[*i, i* + *k* – 1].

A *k-mer order* is a function on *k*-mers *o*: 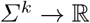. We say that *k*-mer *x*_1_ is *smaller than x*_2_ under *o* (*x*_1_ < *_o_ x*_2_) iff *o*(*x*_1_) < *o*(*x*_2_).

A *de Bruijn graph* (dBG) of order *k* is a directed graph in which every node is labelled with a distinct *k*-mer and there is a directed edge from node *a* to *b* iff the (*k* – 1)-long suffix of *a* is the same as the (*k* – 1)-long prefix of b. The edge is labelled with the (*k* + 1)-long merge of the two labels. A *complete dBG* has a node for every possible *k*-mer and an edge for every possible (*k* + 1)-mer. Paths in a dBG of order *k* represent sequences, and a path of *w* nodes represents a sequence of *w* overlapping *k*-mers.

### 2.2 Minimizers

A *minimizer scheme* is a function *f_k,w,o_*: *Σ*^*w+k*-1^ → {0,…, *w* – 1}. Function *f* returns the position of the minimum *k*-mer under o in a given sequence of *w* overlapping *k*-mers (i.e., in an *L* = *w* + *k* – 1 long sequence). By convention, ties are broken by choosing the left-most *k*-mer. The *minimizers* of a string *S*, denoted as 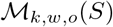, are all the positions in the string that are selected by applying the scheme to all overlapping *L*-long windows of *S*.

#### Minimizer density

The *expected density* of a minimizer scheme is the expected fraction of *k*-mer positions that will be selected as minimizers over an infinitely long random i.i.d. sequence. The *particular density* of a minimizer scheme on a specific sequence *S* (e.g., the human genome) is the fraction of *k*-mer positions selected by the scheme on that sequence.

The *expected* (*particular*) *density factor* is the expected (particular) density multiplied by (*w* + 1).

The expected density factor of random minimizers is 2 (Marçais et al., 2017). Marçais et al. (2018) discuss the asymptotic behaviour of minimizer density as *k* → ∞ or *w* → ∞ and prove a general lower bound of 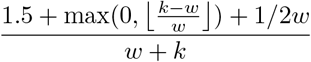 for the density of any forward scheme, generalized sketching schemes of which minimizers are a subset.

#### Partition-compatible minimizer order

Given an ordered partition of *Σ^k^*, 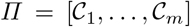, and a *k*-mer order *h*, we say that minimizer order *o_Π,h_* is *compatible with Π* and *h* if for 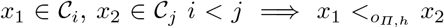 and if *i* = *j* then *x*_1_ < *_o_Π,h__ x*_2_ ⇔ *h*(*x*_1_) < *h*(*x*_2_). In other words, the partition order determines the order between elements from different sets, and another order (typically random) determines the order within sets.

### 2.3 Minimum decycling sets

A *decycling set* of a graph *G* = (*V, E*) is a set of nodes whose removal results in an acyclic graph. Finding an MDS (also called feedback vertex set) in an arbitrary graph is NP-hard (Karp, 1972). We are interested in an MDS of a complete dBG of order *k*. Mykkeltveit (Mykkeltveit, 1972) gave an efficient algorithm to construct such a set, which we denote by 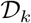, in time linear in the size of a complete dBG, i.e., *Θ*(|*Σ*|*^k^*). A *pure cycle* is a set of nodes corresponding to all the cyclic rotations of some *k*-mer (Mykkeltveit, 1972). Mykkeltveit showed that D_k_ contains a single node from each pure cycle in a complete dBG. Moreover, each pure cycle defines a conjugacy class, and thus the pure cycles factor the complete dBG, namely, every *k*-mer belongs to exactly one of the pure cycles.

#### Mykkeltveit embedding

To determine which of the cyclic rotations of a *k*-mer to include in 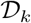, Mykkeltveit defined an embedding of *k*-mers in the complex plane. For a *k*-mer *x*, 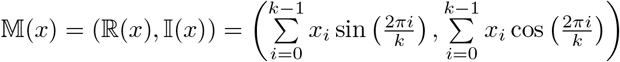 where *x_i_* is the numeric encoding of the *i*-th character of *x* (in our case the encoding of the DNA alphabet is: A=0, C=1, G=2, T=3) (Figure 1). We say a rotation *x* is *positive* if 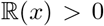. All positive rotations of a *k*-mer are consecutive. The MDS constructed by Mykkeltveit’s algorithm includes, for each conjugacy class, the first positive counter-clockwise rotation. When all rotations have 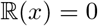, any arbitrary *k*-mer from the cycle can be selected.

**Fig. 1:**
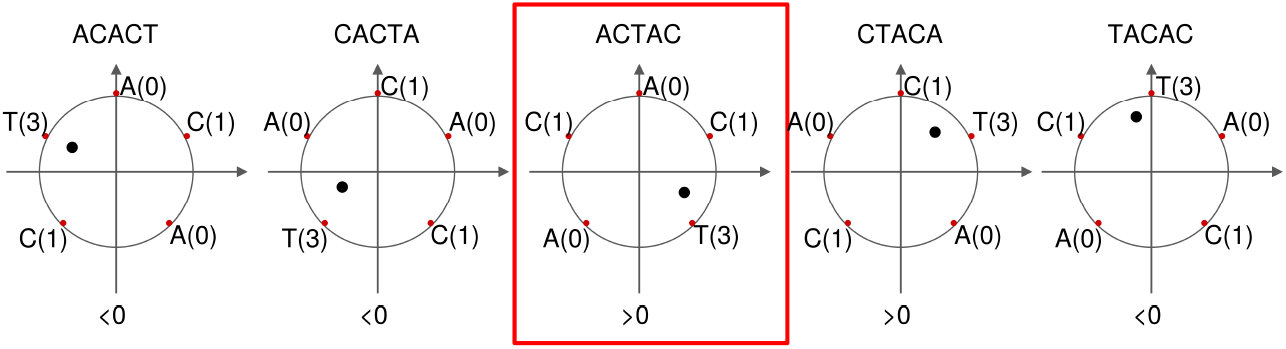
Mykkeltveit embedding. The embedding is shown for the rotations of the *k*-mer ACACT, indicated above each subfigure. Each letter of the *k*-mer corresponds to a weight (in parentheses) placed at the *k*-th roots of unity (red dots). The embedding represents the center of mass of the *k*-mer (black dot). The sign of each embedding projected onto the real axis is shown below each rotation. In this example, ACTAC (red box) is the first counter-clockwise rotation *x* with 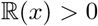, and is thus included by Mykkeltveit’s algorithm in the MDS.

Mykkeltviet’s algorithm has an efficient implementation due to Knuth (Knuth, 2003). This implementation uses the FKM algorithm (Fredricksen and Maiorana, 1978) to enumerate the *k*-mer conjugacy classes in lexicographic order. The representative selected for each class is the first positive one, and for classes with 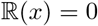 for all *k*-mer rotations, the lexicographically smallest *k*-mer is included in the decycling set. An MDS consists of *Θ*(|*Σ*|*^k^*/*k*) *k*-mers.

### 2.4 Universal hitting sets

A *universal hitting set* (UHS) 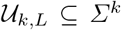 is a set of *k*-mers such that any *L*-long string contains at least one *k*-mer from 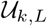 as a contiguous substring.

By construction, at least one *k*-mer from 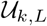 must appear in every window of *w* = *L* – *k* +1 overlapping *k*-mers, and thus the nodes represented by 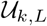 will be a covering set for all w-long paths in a complete dBG of order *k* (Marçais et al., 2017).

Two algorithms were proposed for UHS construction, DOCKS (Orenstein et al., 2017) and PASHA (Ekim et al., 2020). They first generate an MDS using Mykkeltveit’s algorithm. Both algorithms then add *k*-mers greedily until no path of length *w* remains in the graph. DOCKS uses dynamic programming to compute the number of w-long paths in the remaining graph, resulting in a runtime of *O*((1 + *p*)|∑|^*k*+1^*L*) for p iterations of node removal. Using DOCKS, small UHSs have been constructed only for *k* ≤ 11. Minimizer orders compatible with these UHSs were shown to have lower density than random orders (Marçais et al., 2017).

PASHA uses a randomized approximation algorithm for Set Cover to remove a small number of nodes in the remaining graph, resulting in a runtime of *O*((*L*^2^|*Σ*|^*k*+1^*log*^2^(|*Σ*|^*k*^))/(*ϵδ*^3^)) where *δ* and *ϵ* are parameters of the approximation guarantee. UHSs have been constructed using PASHA only for *k* ≤ 13 and the density of minimizer orders compatible with them was observed to be slightly higher than that of minimizer orders compatible with DOCKS UHSs.

Miniception (Zheng et al., 2020b) constructs a UHS for parameters *k* and *L* with an additional parameter *k*_0_ < *k*. *k*-mers with the first or the last *k*_0_-mer as their minimizers are added into the UHS. This set does not need to be pre-computed, and instead membership in the set can be determined on the fly for each *k*-mer in a sequence. Minimizer orders compatible with Miniception-generated UHSs can therefore be computed efficiently for any *k*, but were shown to have higher density than orders compatible with UHSs constructed by PASHA.

## 3 Methods

### 3.1 Decycling-set-based minimizer orders

Given an MDS 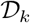, we define a *k*-mer order in which *k*-mers in 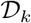 precede all other *k*-mers, and for pairs not determined by this rule the order is random. Formally, we define the partition 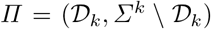 and a pseudo-random *k*-mer order *h*, and use a minimizer order compatible with Π and h. We implement a pseudo-random order efficiently by XOR-ing the binary representation of a *k*-mer with a random 2*k* bit seed as was done in (Wood and Salzberg, 2014; Flomin et al., 2022). 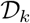 can be constructed efficiently using Knuth’s implementation of Mykkeltveit’s algorithm (Mykkeltveit, 1972) as described above.

For large values of *k*, when 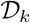 is too large to store or takes too much time to compute, we instead scan the target sequence and, for every *k*-mer test its membership in 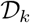 on the fly using the procedure outlined in Algorithm 1 (Figure 2). The real parts of the embeddings of a *k*-mer *x* and its clockwise rotation x’ are computed in *O*(*k*) time and compared to determine if *x* is the first positive counter-clockwise rotation. If 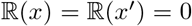, then the algorithm determines whether *x* is a lexicographically smallest rotation in *O*(*k*) time.

**Fig. 2:**
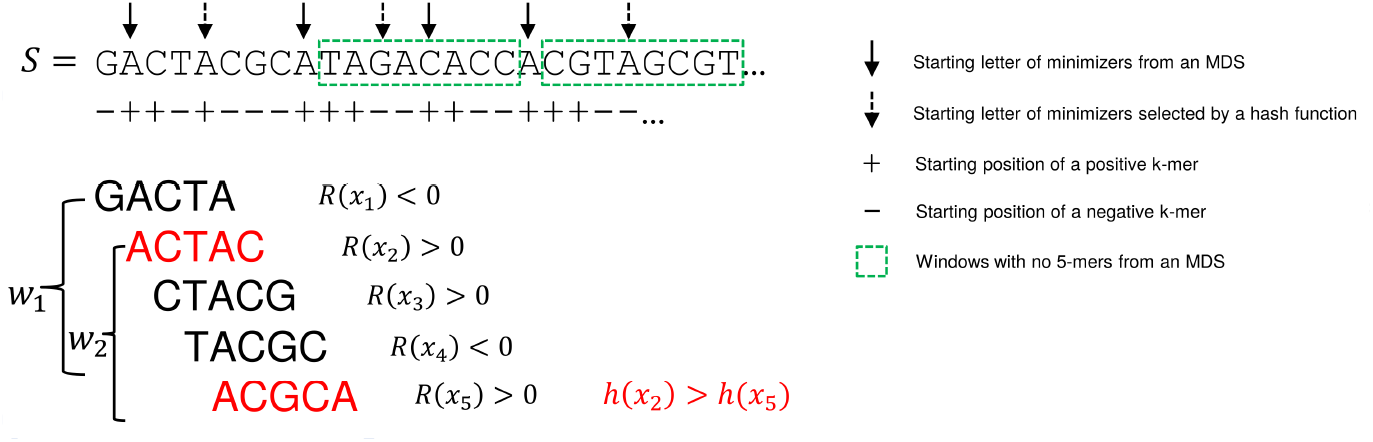
Decycling set compatible minimizers. An example of selecting minimizers based on an MDS with parameters *k* = 5, *L* = 8. *k*-mers in the leftmost two windows are shown below the sequence, with *k*-mers in the decycling set colored red. The second window *w*_2_ contains two *k*-mers from the decycling set, and a hash function was used to select ACGCA as the minimizer. Sequences boxed in green represent windows with no *k*-mer from the decycling set, and thus a random *k*-mer was selected as the minimizer by a hash function.

#### Algorithm 1

Decycling-set membership

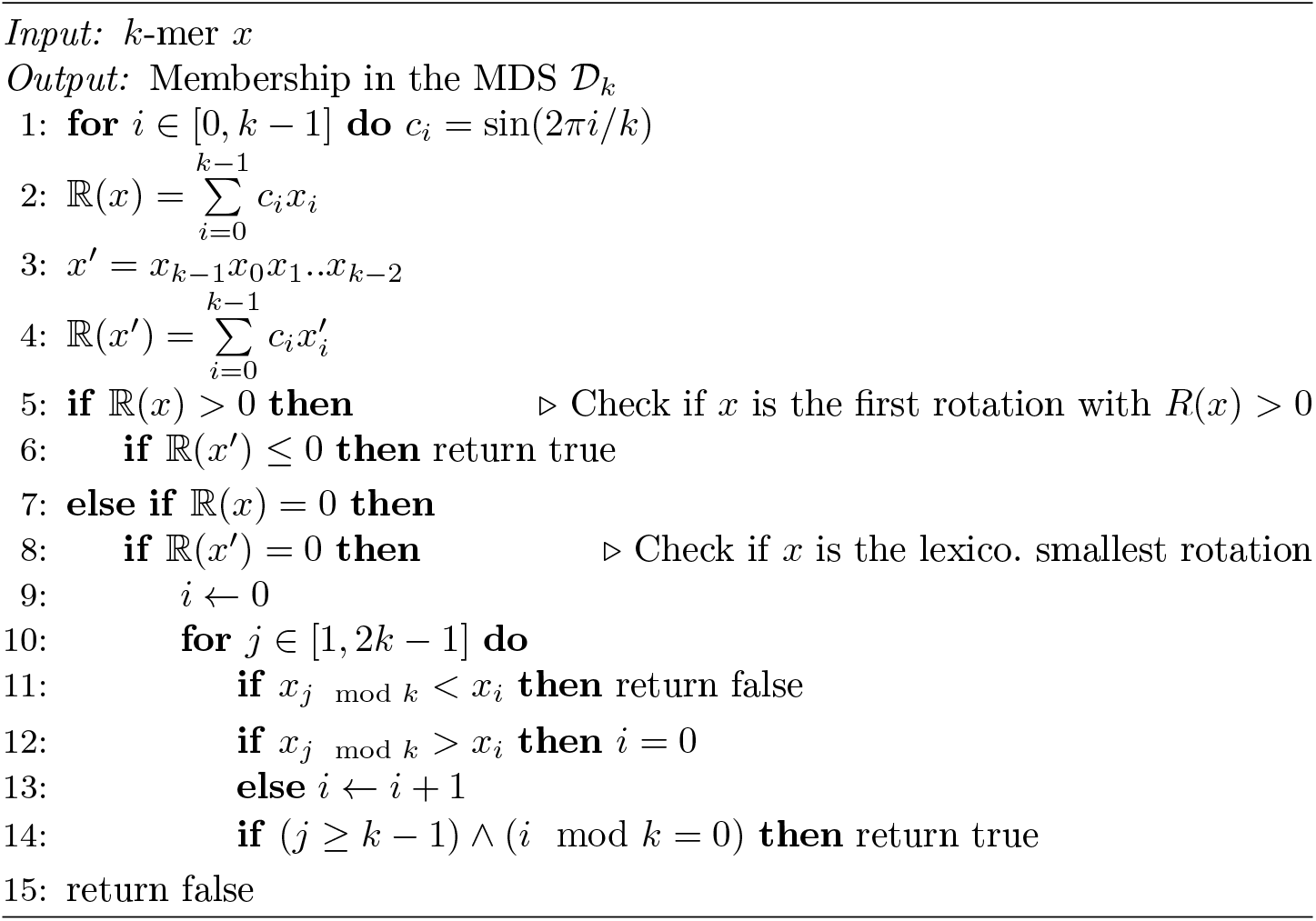

#### Proposition 1 (Alg. 1 correctness).

Alg. 1 correctly determines whether a *k*-mer is a member of 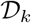 in time *O*(*k*).

*Proof*. The proof follows from the definition of 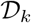. We say that a *k*-mer *x* is *positive, negative*, or *non-positive* if 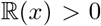, < 0, or ≤ 0, respectively. Recall that a *k*-mer 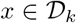 iff either: (i) it is the first positive counter-clockwise rotation in its conjugacy class; or, (ii) all *k*-mers in the conjugacy class have 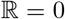 and *x* is a lexicographically smallest rotation.

For (i), line 6 returns true iff the input *k*-mer *x* is the first positive counterclockwise rotation in its conjugacy class, i.e., *x* has 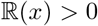 and the one-letter clockwise rotation of *x*, denoted *x’*, has 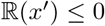.

For (ii), note that if two consecutive rotations of a *k*-mer *x, x’* have 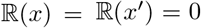 (lines 7-8), then all *k*-mers in that conjugacy class have zero embedding (Lemma 1 in Mykkeltveit (Mykkeltveit, 1972)). The loop in lines 10–14 checks all possible rotations of *x* and returns false if it finds a *k*-mer that is lexicographically smaller than *x* (line 11). Otherwise it returns true if either it checked all possible rotations and none of them is lexicographically smaller than *x* (i = 0 and *j* = *k* – 1) or it found that *x* is identical to one of its rotations and *x* is a lexicographically smallest rotation (*i* = *k* and *j* ≥ *k* – 1) (line 4).

The embedding computations (lines 1, 2, and 4) take *O*(*k*) time. The loop beginning on line 10 can run for at most 2*k* times and performs constant time computations per iteration for a total running time of *O*(*k*).

### 3.2 Double decycling-set-based minimizer orders

By symmetry, Mykkeltveit’s construction can be used to create an MDS using the first counter-clockwise negative *k*-mer *x* in each conjugacy class rather than the first positive one. We refer to this set as the *symmetric* decycling set 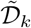. The decycling set and symmetric decycling set divide sequences according to the following interesting property:

#### Theorem 1 (remaining path partition).

*In any remaining path in a complete dBG after removing 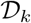, all the positive nodes (i.e., labelling positive *k*-mers) precede all the non-positive nodes*.

In other words, a remaining path must consist of two distinct parts: A *positive part*, containing only positive *k*-mers, followed by the *non-positive part* consisting of only non-positive *k*-mers. The proof relies on two lemmas:

#### Lemma 1.

*The k-mers associated with all incoming neighbours of a node x in a dBG have the same* 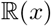.

*Proof*. All incoming neighbours *y* of *x* differ only in *y*_0_, and have embedding with 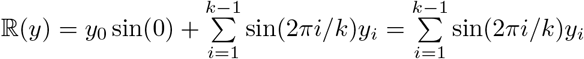.

#### Lemma 2.

*The pure cycles factor a complete dBG, namely, every k-mer belongs to exactly one of the pure cycles*.

*Proof*. Every *k*-mer is on some pure cycle corresponding to its rotations. Assume the contrary that *k*-mer *x* is on two distinct pure cycles, *C*_1_ and *C*_2_. Let y be the last common node in the path in *C*_1_ ∩ *C*_2_ starting from x. Then, the edges out of y in the two cycles are distinct, contradicting the fact that both correspond to the cyclic rotation of y.

*Proof* (*Thm. 1*). Let *x_i_* be the first non-positive node in a remaining path *x*_1_,…, *x_t_* and assume the contrary that there exists a positive *x_j_* for *j* > *i*. W.l.o.g. assume *x_j_* is the first with that property in the path. Let C be the pure cycle that contains *x_j_*. *C* exists and it is well defined by Lemma 2. Let *y* be the node preceding *x_j_* in *C*. By Lemma 1, 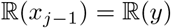. Since *y* is non-positive, *x_j_* should be in 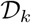 as the first positive node in *C*, a contradiction.

By a similar argument, in a remaining path after removing 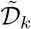, the negative nodes precede all other nodes. Thus, removal of a *double decycling set* consisting of 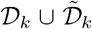 would leave only short remaining paths that cannot contain both negative and positive *k*-mers.

We define a partition-compatible minimizer order based on double decycling sets with 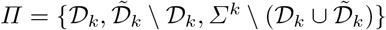. Because the double decycling set leaves even shorter remaining paths, we hypothesize that this minimizer order will achieve lower density compared to the one using only a single decycling set.

## 4 Results

We compared the performance of our new MDS-based minimizer orders to random orders, and orders based on DOCKS, PASHA, and Miniception (Zheng et al., 2020b), across a range of *k* and *L* values. We measured performance using expected and particular density factors. Expected density factors were estimated by measuring density on ten random i.i.d. sequences of 10M nt, each time with a different pseudo-random seed for the *k*-mer order. Particular density factors were measured on ten randomly selected 10M nt segments from chromosome *X* of the CHM13 telomere-to-telomere human genome assembly (Nurk et al., 2022) using different pseudo-random seeds. Miniception uses lexicographic order by default, which was shown to have worse performance than random order. To be fair, here we modified Miniception to use the same random hash as our implementations and ran it with the recommended parameters. Results of further experiments on more particular genome sequences can be found in the Supplement. Our code for implementing the different minimizer orders is publicly available at github.com/OrensteinLab/DecyclingSetBasedMinimizerOrder.

### 4.1 MDS-based orders outperform UHS-based orders

Figure 3 compares the density factors of the tested orders for *k* = 11 and 10 ≤ *L* ≤ 200, and for 5 ≤ *k* ≤ 15 and *L* = 100. Average density factors over the repeated runs are shown for visual clarity. The same plots with error bars and theoretical lower bounds displayed are available in Supplementary Figure S1.

**Fig. 3:**
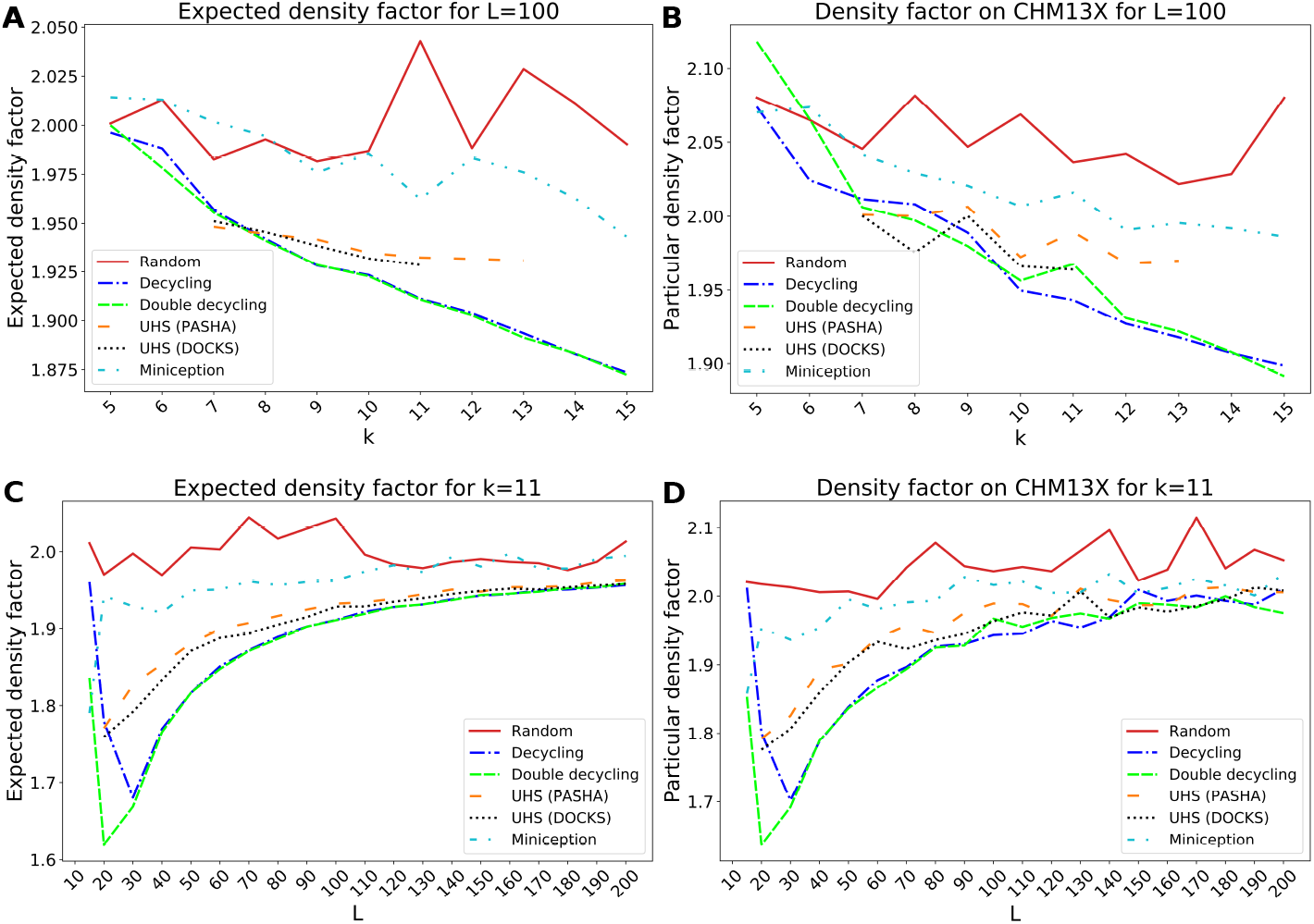
Density factor of MDS-based and UHS-based orders. The expected density (**A,C**) and particular density factors on CHM13X (**B,D**) of different minimizer orders is compared for *L* = 100 and varying 5 ≤ *k* ≤ 15 (**A,B**) and for *k* = 11 and 10 ≤ *L* ≤ 100 (**C,D**).

UHSs for *k* < 7 and *L* = 100 consist only of an MDS, and thus are not shown. DOCKS-generated UHSs were computed up to *k* = 11 and PASHA-generated UHSs up to *k* = 13. UHSs for larger *k* could not be generated due to the runtime and storage required for every combination of *k* and L. In contrast, our new MDS-based orders have the distinct advantage of being easily computable on the fly for any (k, L) combination.

The MDS-based orders consistently perform similarly or better than UHS-based orders. Miniception performs better than random and worse than the UHS-based orders, as was previously shown (Zheng et al., 2020b). As expected, random orders typically perform worst across most of the range of L, and the relative improvement of UHS- and MDS-based orders compared to random orders increases with *k*. Conversely, as *L* grows for fixed *k* the density factors of the different methods tend to perform similar to random since longer windows are more likely to contain multiple *k*-mers from the sets defining the order (UHS or MDS), and the *k*-mers within the set are ordered randomly. MDS-based orders have much lower standard errors than the others, likely because the decycling set *k*-mers remain the same for all repetitions regardless of the random seed.

The particular density factor is slightly higher than the expected density factor and is more variable, with higher standard errors, as it is dependent on the particular sequence, and *k*-mer usage is not uniformly distributed in real genomes. However, the overall shape of the density curves and the performance ranking among the methods remain the same. Particular densities for more sequences are presented in Figure S2.

### 4.2 Scaling MDS-based orders to *k* ≥ 20

We compared the MDS-based orders to the random baseline order and to Miniception for values of *k* greater than feasible to run with DOCKS and PASHA. Figure 4 shows results for *k* = 20, 50, and 100. Average density factors over the repeated runs are shown. The same plots with error bars displayed are available in Supplementary Figure S1 and particular densities for more sequences are presented in Figure S2.

**Fig. 4:**
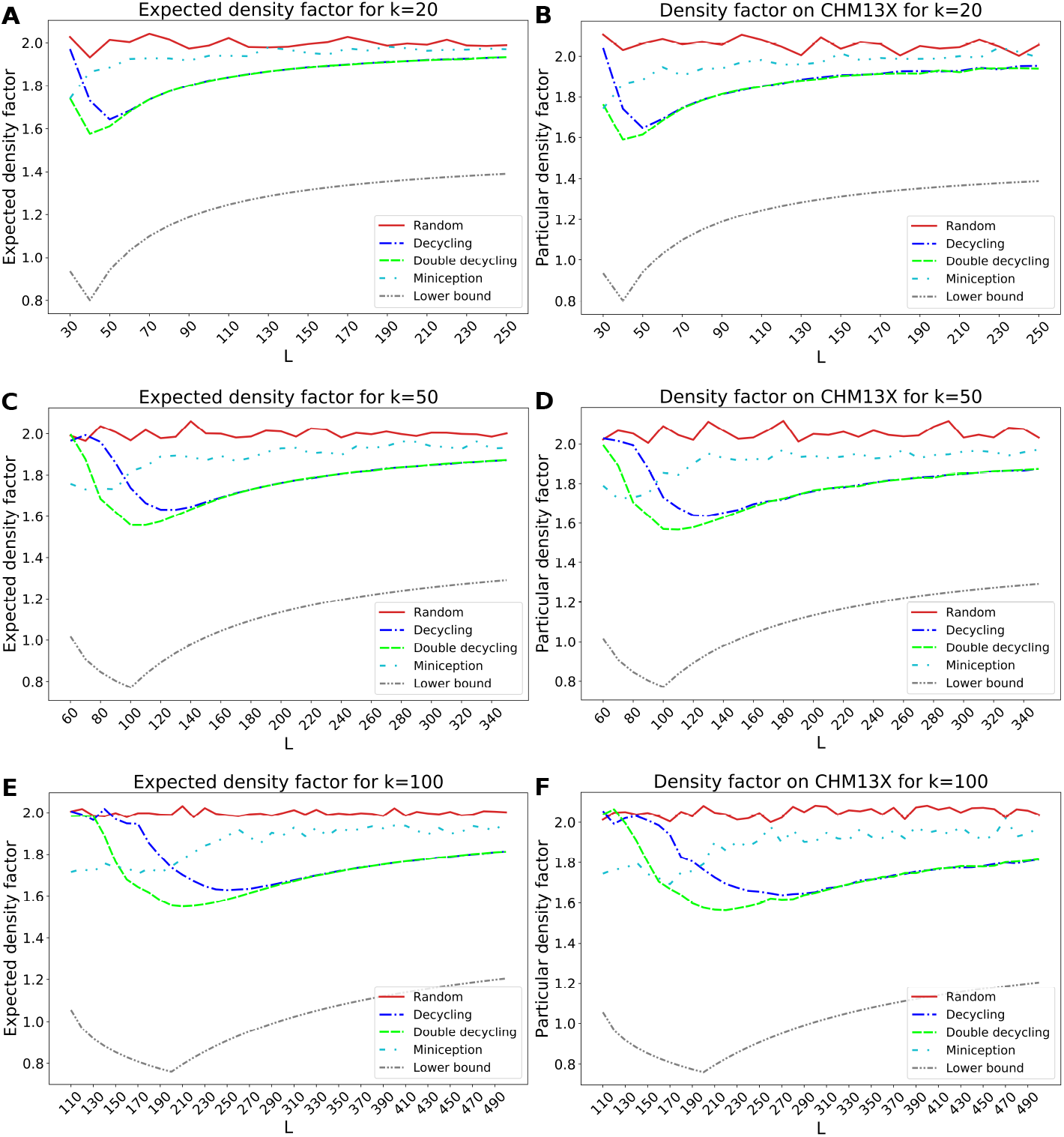
Density factors of MDS-based orders for large *k*. The expected density factor (**A,C,E**) and the particular density factor on CHM13X (**B,D,F**) of different minimizer orders is compared for *k* = 20 (**A,B**), *k* = 50 (**C,D**), and *k* = 100 (**E,F**) over variable *L* values.

As *k* grows, the advantage of the MDS-based orders becomes even more pronounced and the double decycling set-based order improves more significantly over the decycling-set-based order. This is true in particular for smaller *L*, with the differences between the decycling- and double decycling-set-based orders beginning to diminish as *L* > 2*k*. The advantage compared to Miniception is maintained for large values of *k* across most *L* values, and the MDS-based minimizer orders achieve density factors that are up to 20% lower than Miniception-based orders. Miniception-based orders achieve lower density for small *L* as MDS-based orders are similar to random in that regime, since most windows do not contain a member of the MDS. In contrast, Miniception approaches its theoretical density factor of 1.67 while *L* ≤ 2*k* and converges to (but remains below) the performance of random only for larger L. In most cases, the particular density factor is slightly higher than the expected density factor.

There exists a consistent gap between the theoretical lower bound for forward schemes, which is not known to be tight for minimizers, and converges to 1.5 as *L* increases, while the MDS-based orders converge to 2. Interestingly, the shape of the density factor curves is similar with a minimum around *L* = 2*k*.

### 4.3 Runtime and memory usage

We report runtime measurements under three regimes of our decycling-set-compatible minimizer scheme implementations. For *k* < 20, the MDS is pre-computed and stored as a boolean vector. Thus, *k*-mer set membership is determined with an O(1) lookup. The exponential growth of the MDS makes it impossible to pre-compute or store in memory for larger *k*. Instead, we compute *k*-mer set membership on the fly for every *k*-mer in *O*(*k*) time using Algorithm 1. As *k*-mers are represented using two bits per nucleotide, our implementation in C++ can use CPU-supported 128-bit operations for *k* ≤ 63 (an additional bit of the representation is used to indicate set membership). We used the GMP library (https://gmplib.org/) to support operations for *k* > 63. As a result, the process is an order of magnitude slower than CPU-supported operations. We report runtimes to compute minimizers of a random 10M nt sequence, for all methods and three values of *k* representing the different regimes in Table 1. Runtimes were measured on a 44-core, 2.2 GHz server with 792 GB of RAM, using a single thread, and are averaged across ten runs.

**Table 1:**
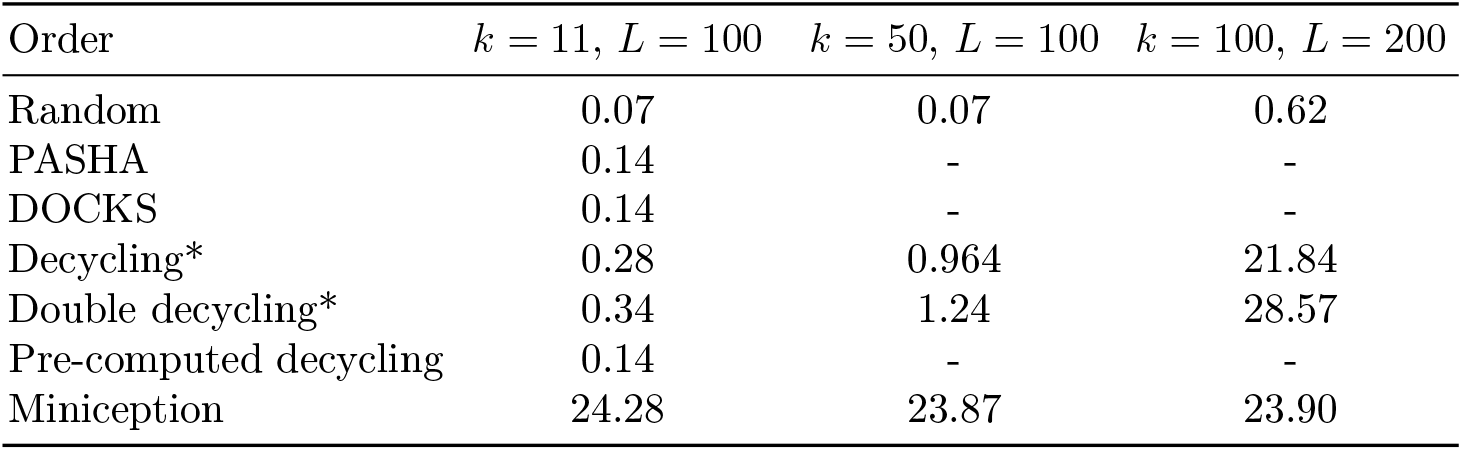
Runtime results. Average time in seconds to compute the minimizers for each order on a 10M nt sequence. *MDS membership is computed on the fly.

| Order | $k = 11, L = 100$ | $k = 50, L = 100$ | $k = 100, L = 200$ |
| --- | --- | --- | --- |
| Random | 0.07 | 0.07 | 0.62 |
| PASHA | 0.14 | - | - |
| DOCKS | 0.14 | - | - |
| Decycling* | 0.28 | 0.964 | 21.84 |
| Double decycling* | 0.34 | 1.24 | 28.57 |
| Pre-computed decycling | 0.14 | - | - |
| Miniception | 24.28 | 23.87 | 23.90 |

We implemented the pseudo-random order by a simple XOR operation and a comparison of the resulting integers. Thus, the random order is extremely fast for all *k* and *L* values. The lookup and comparison used by the pre-computed decycling set and UHS methods is also fast, but limited to relatively small *k*. Algorithm 1 with the more complex embedding calculation and comparison is slower and the runtime increases with *k*. Our implementation’s runtime increases by 1-2 orders of magnitude for *k* > 63. However, even the slowest double-decycling order processes 10M nt in less than half a minute for *k* = 100, achieving much lower density than the random order. Miniception is implemented in Python and has a consistent runtime across all values of *k*, but is much slower than other methods in two of the three regimes.

Table 2 presents the memory usage for the pre-computed MDS-based order for *k* = 5, 10, 15, 18 over *L* = 100 for a 10M nt sequence measured as the maximum resident set size (RSS). For the lookup-based methods, which store a pre-computed set, memory grows exponentially with *k*, and, for a 10M nt sequence, the memory needed to store the set begins to dominate the memory for the sequence starting at *k* = 14. The required memory becomes infeasible prohibitively large for *k* = 20. The memory overhead of the random and MDS-based orders computed on the fly is negligible, and so the memory usage is dominated by the sequence being processed.

**Table 2:** Maximum memory usage. Peak RAM usage of the lookup-based pre-computed MDS-based order for *L* = 100 on a 10M nt sequence.

| | $k = 5$ | $k = 10$ | $k = 15$ | $k = 18$ |
| --- | --- | --- | --- | --- |
| Maximum memory usage | 16.5MB | 16.7MB | 147.7MB | 8.4GB |

## 5 Discussion

In this study we developed a method to generate highly effective minimizer orders for any *k*. A major limitation of minimizer orders based on UHSs was the need to create and store the whole set in advance. Instead, we based the order only on the MDS, avoiding the need to add *k*-mers to make the set universal. This bypasses the most costly steps in DOCKS and PASHA, and generates minimizer orders that are even better in terms of their density factors. Furthermore, this approach enables calculating the minimizers in a sequence efficiently on the fly, without the need to store the set. We showed that based on Mykkveltveit’s algorithm, we can determine in *O*(*k*) if a *k*-mer belongs to the MDS, and thus MDS membership can be checked for all *k*-mers in a sequence. The resulting new orders are comparable or better in their density than UHS-based minimizer orders, thus achieving good performance while avoiding escalating runtime and memory usage with the increase of *k*.

In addition, we defined double decycling-set-based orders. For larger *k* and *L* ≤ 2*k*, the double decycling-set-based orders yielded much lower density than the decycling-set-based orders (Figure 3) at the cost of a small increase in runtime. As the density of the two methods becomes similar as *L* increases, we recommend using double decycling-set-based orders for *L* ≤ 2.5*k* to achieve lower density, while decycling-set-based orders can be used for *L* > 2.5*k* to achieve similar density with slightly faster runtime. When *L* ≫ *k*, our new MDS-based orders achieved only a modest advantage in density over the much faster random order(Figure 4).

We see several promising future directions. Our work focused on general minimizer orders, but other sequence sketches are sequence-specific or waive the strict window-guarantee of minimizers to obtain improved performance. The advantages of an MDS are likely to extend to these methods. For example, frequency-based orders are known to be highly efficient in terms of density and are easily computable as sequence-specific minimizer orders. It will be interesting to extend our work by ranking each of the sets in a partition by their frequency in a specific sequence dataset to achieve lower density values (as was recently shown by incorporating UHS-based orders with frequency ranking (Nyström-Persson et al., 2021)). In addition, it would be possible to use decycling sets and their variants as sketches without defining compatible minimizer orders by simply including all decycling set *k*-mers in the sketch. While not providing a window guarantee, such schemes would be better conserved under mutations than minimizers, as they are not dependent on a longer sequence window (Edgar, 2021).

There are also open theoretical and technical questions arising from our study. It is necessary to explain why the densities of the MDS-based minimizer orders match the shape of the theoretical lower bound for the more general forward sequence sketching schemes. The gapobserved between the MDS-based order and the lower bound should be studied, and hopefully a tighter lower bound for minimizer schemes can be found. Performance analysis of our new minimizer orders in terms of other criteria, including maximum bin load (Chikhi et al., 2016; Flomin et al., 2022) and conservation (Edgar, 2021), is also a worthy goal. Immediate future work can improve the implementation for *k* > 63 to speed up runtimes for very large *k*.

To conclude, our new approach can enable more efficient analyses of high-throughput sequencing data. By implementing our new MDS-based minimizer orders in data structures and algorithms of high-throughput DNA sequencing analysis, we expect to achieve reductions in runtime and memory, beyond what was previously demonstrated using UHS-based minimizer orders.

## Acknowledgments

This study was supported by a United States–Israel Binational Science Foundation (BSF) grant no. 2020297 to YO and BB. RS was supported in part by the Israel Science Foundation (grant 2206/22) and by Len Blavatnik and the Blavatnik Family foundation. DP and LP were supported in part by fellowships from the Edmond J. Safra Center for Bioinformatics at Tel-Aviv University. LP was supported in part by the National Natural Science Foundation of China project 61902072. BE and BB were partially supported by grant NIH 1R35GM141861 (to BB).

