## Supplementary File for "Efficient minimizer orders for large values of *k* using minimum decycling sets"

Supplementary results are shown in Supplementary Figures S1 and S2.

* shared correspondence

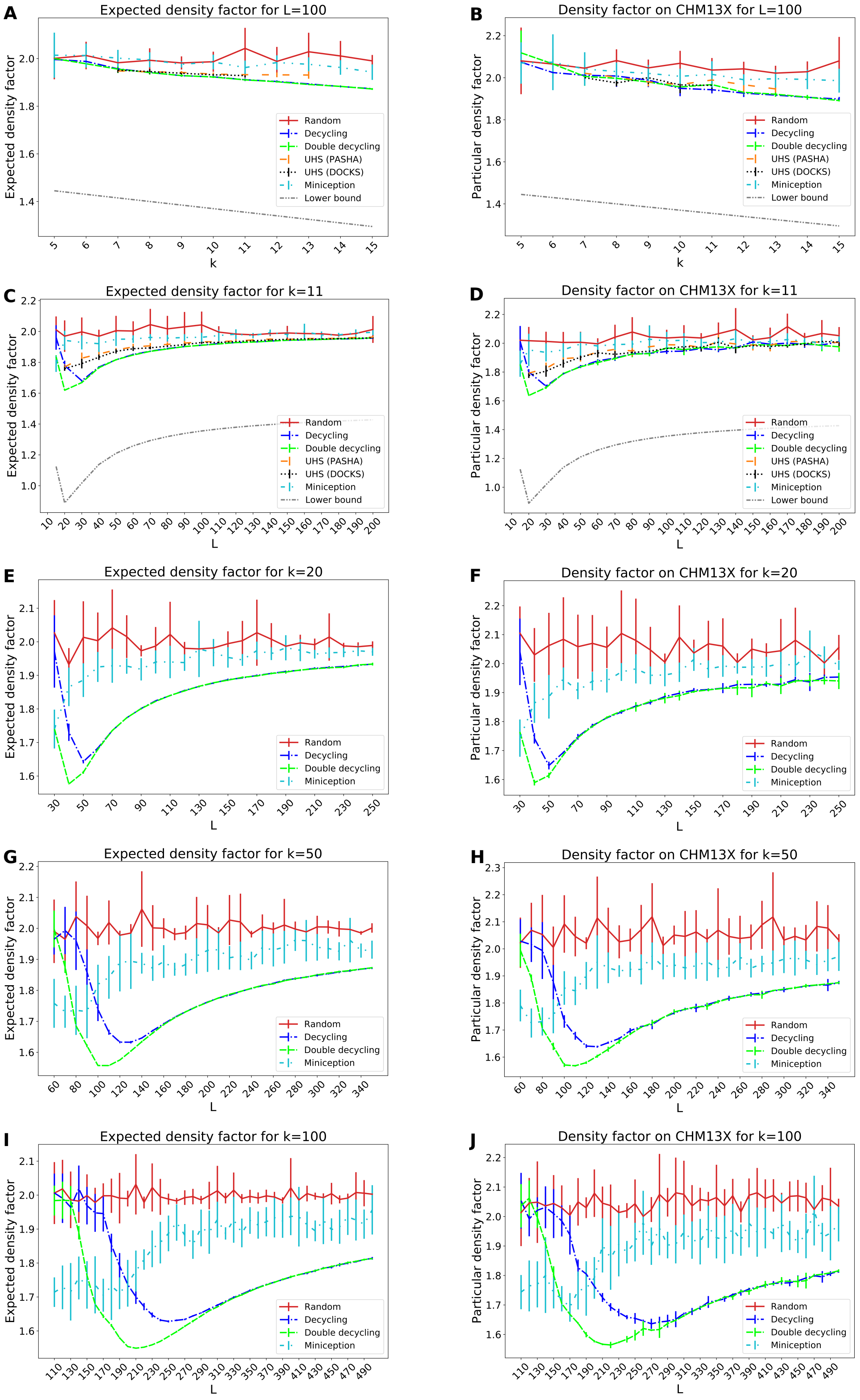


Fig. S1: **Density factor for all methods including error bars.** The expected density (left) and particular density factors on samples from CHM13X (right) of different minimizer orders is compared over $L = 100$ and $5 \leq k \leq 15$ (**A**,**B**) and for a range of fixed $k$ with varying $L$ (**C**-**J**). The averages and standard errors over 10 runs with 10M nt sequences are shown.

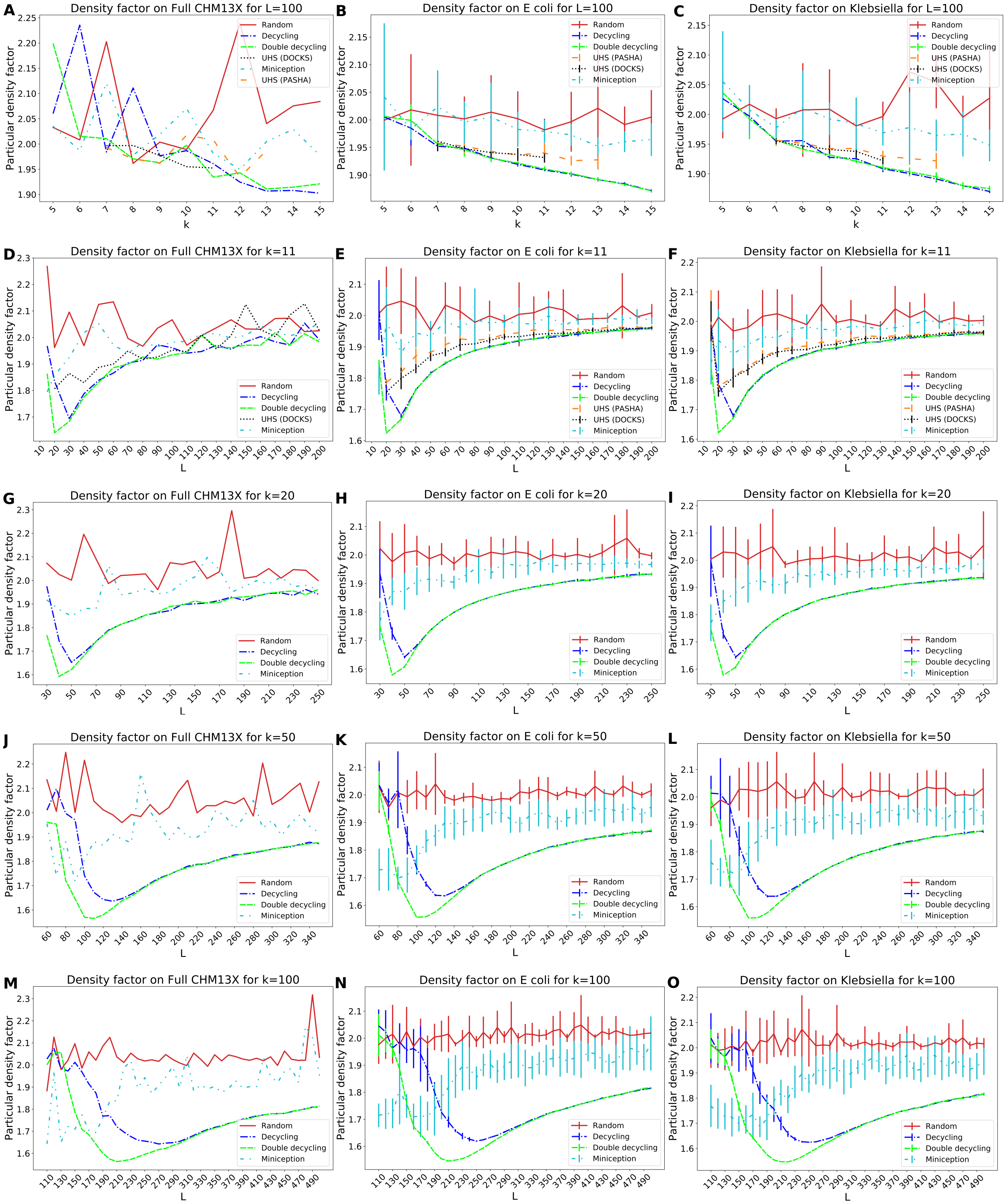

Fig. S2: **Particular density factors various sequences including error bars.** The particular density factors are shown for: the entire 154M nt sequence of chromosome X from CHM13 (**left**); the 4.9M nt genome of *E coli* strain W (RefSeq accession GCF_000184185.1) (**center**); and, the 5.3M nt genome of *Klebsiella pneumoniae* strain HS11286 (RefSeq accession GCF_000240185.1) (**right**). For the bacterial genomes, averages and standard errors over 10 runs are shown.
